# Primary human myoblasts display only minor alternative polyadenylation compared to the transformed C_2_C_12_ model of muscle differentiation

**DOI:** 10.1101/2023.12.17.572066

**Authors:** Akriti Varshney, Paul F. Harrison, Angavai Swaminathan, Sarah E. Alexander, Bernhard Dichtl, Séverine Lamon, Traude H. Beilharz

**Affiliations:** Development and Stem Cells Program, Monash Biomedicine Discovery Institute and Department of Biochemistry and Molecular Biology, Monash University, Melbourne, VIC 3800, Australia; Institute for Physical Activity and Nutrition, School of Exercise and Nutrition Sciences, Deakin University, Geelong, Victoria 3220, Australia; School of Life and Environmental Sciences, Deakin University, Geelong, Victoria 3220, Australia; Monash Bioinformatics Platform, Monash University, Melbourne, VIC 3800, Australia

**Author notes:** Traude Beilharz, Development and Stem Cells Program, Monash Biomedicine Discovery Institute and Department of Biochemistry and Molecular Biology, Monash University, Melbourne, VIC 3800, Australia.

**Keywords:** alternative polyadenylation, cholesterol biosynthesis, primary human myoblast differentiation

## Abstract

Alternative polyadenylation has been linked to multiple developmental and disease transitions. The prevailing hypothesis being that differentiated cells use longer 3’ UTRs with expended regulatory capacity whereas undifferentiated cells use shorter 3’ UTRs. Here, we describe the gene expression and alternative polyadenylation profiles of human primary myoblasts over a time course of differentiation. Contrary to expectations, only minor changes in the 3’ end choice were observed. To reconcile this finding with published research, we devised a new bioinformatic method to compare the degree of alternative polyadenylation in the differentiation of primary human and immortalized murine (C_2_C_12_) myoblasts. Differentiated human primary myotubes display only half the alternative polyadenylation of the mouse model, with less than 1/10 of the genes undergoing alternative polyadenylation in C_2_C_12_ cells showing evidence of alternative processing in human primary muscle differentiation. A global reduction in the expression of cleavage and polyadenylation factors in C_2_C_12_, but not in primary human myotubes may explain the lack of alternative polyadenylation in this system. Looking more broadly at transcriptome changes across differentiation shows that less than half of the genes differentially expressed in the immortalized model were recapitulated in primary cells. Of these, important metabolic pathways, such as glycolysis and sterol biosynthesis, showed divergent regulation. Collectively, our data caution against using immortalized cell lines, which may not fully recapitulate human muscle development, and suggest that alternative polyadenylation in the differentiation of primary cells might be less pronounced than previously thought.

## Introduction

Alternative polyadenylation (APA) is a co-transcriptional process that generates mRNA isoforms with different 3’ untranslated regions (UTRs) from the same genetic locus. The cleavage and polyadenylation (CPA) machinery responsible for APA consists of several evolutionarily conserved protein complexes, including cleavage and polyadenylation specificity factor (CPSF), cleavage stimulation factor (CstF), Cleavage factor I and II (CFI and CFII, and a poly(A) polymerase (Giammartino et al. 2011; Tian and Manley 2017). The CPSF component is primarily responsible for detecting a canonical AAUAAA poly(A) signal (PAS) in the 3’ UTR, or related non-canonical PAS for downstream endonucleolytic pre-mRNA cleavage and the non-templated addition of adenosine nucleotides for poly(A) tail synthesis (Giammartino et al. 2011; Tian and Manley 2017). When more than one poly(A) signal exists, the choice is regulated by multiple intersecting mechanisms, including the strength of PAS and the relative concentration of the cleavage and polyadenylation machinery (Gruber et al. 2014). The length of the 3’ UTR is significant because it can contain regulatory elements for nuclear export, stability, and spatiotemporal translation of mature mRNA (Turner et al. 2018; Beilharz et al. 2019; Pereira-Castro and Moreira 2021). Seminal early work showed that 3’ UTR shortening resulted in aberrant expression of key oncogenic genes by loss of crucial RNA-binding protein and microRNA binding sequences (Sandberg et al. 2008; Mayr and Bartel 2009). The use of proximal PAS, and thus shorter 3’ UTRs, has now been associated with proliferative growth in multiple stem cell and cancer models (Kandhari et al. 2021; Yuan et al. 2021).

The differential expression of CPA factors can explain the relationship between proliferation and APA (Gruber et al. 2014). For example, altered expression of CFIm25 and PCF11 has been linked to 3’ UTR shortening and tumorigenesis in glioblastoma (Masamha et al. 2014) and neuroblastoma (Ogorodnikov et al. 2018). In contrast, distal poly(A) site selection and longer 3’ UTRs have been widely reported in cell differentiation (Ji et al. 2009; Grassi et al. 2019; Sommerkamp et al. 2021). For example, high levels of the polyadenylation factor Fip1 promote the shortening of 3’ UTRs essential for embryonic stem cell maintenance and proliferation (Lackford et al. 2014). Similarly, the differentiation of immortalised skeletal myoblasts leads to the loss of PCF11 with global 3’ UTR lengthening (Wang et al. 2019). APA has also been reported in multiple models of cardiac myotube differentiation and disease (Ji and Tian 2009; Nimura et al. 2016; Soetanto et al. 2016; Yang et al. 2022).

Skeletal muscle development, or myogenesis, is a complex and highly regulated cellular process and has been the model of choice for investigating the role and regulation of APA in cell differentiation (Li et al. 2015; Wang et al. 2019). The model relies on a population of quiescent muscle stem cells, termed myoblasts or satellite cells, that exit their proliferating state and undergo terminal differentiation (Bentzinger et al. 2012). Differentiating muscle cells or myotubes then fuse to become multinucleated myofibers (Chal and Pourquié 2017; Jiwlawat et al. 2018), the contractile units that allow for muscle movement in adults (Korthuis 2011). Some myoblasts retain their stem properties in adult tissues, allowing postnatal skeletal muscle growth and regeneration (Charge and Rudnicki 2004). One of the most common models of mammalian muscle development used for transcriptomic and metabolic studies is the immortalised murine C_2_C_12_ cell line (Burattini et al. 2004).

The myoblast-derived C_2_C_12_ line displays cycles of ‘active’ to ‘resting’ that closely resemble those of satellite cells that cycle between activated and quiescent states (Yoshida et al. 1998). In contrast, primary muscle cell lines are directly established from mouse tissue or human muscle biopsies (Zacharewicz et al. 2020). Although more complex to establish and expensive to maintain, these provide an alternative model that may be molecularly and metabolically more relevant to the human physiology (Abdelmoez et al. 2020).

Here, we sought to understand the extent to which APA is associated with human myoblast differentiation. We compared the transcriptomic and APA profiles of differentiating primary human skeletal myoblasts (*pHSkMs*) with public data from C_2_C_12_ myotubes (Abdelmoez et al. 2020). In contrast to our hypothesis, and unlike what was observed in C_2_C_12_ cells, little APA and only a subtle trend toward 3’ UTR lengthening was found in differentiating *pHSkM*. Differences in proliferative capacity, and stable expression of CPA factors, when compared to C_2_C_12_ cell differentiation, where their expression is globally repressed, may explain the lack of APA in differentiated *pHSkM*. In addition, the models displayed considerable differences in metabolic gene ontology (GO) pathway enrichment, for example, a significant induction of cholesterol biosynthesis in *pHSkM* cells was not recapitulated in C_2_C_12_ myotubes. Collectively, our data suggest caution should be exercised when using immortalized models to infer human skeletal muscle biology and that the connection between APA and differentiation might be more context-dependent than previously thought.

## Results

To investigate APA in human skeletal muscle differentiation, *pHSkM* from two healthy donors, male (22.9 years old) and female (26.6 years old), were established, passaged, and differentiated according to our previously published method (Lamon et al. 2014; Zacharewicz et al. 2020). Samples were collected from the male-derived HSM1 and female-derived HSM2 *pHSkM* lines, grown in replicates and harvested at indicated time points (Fig. 1A). For comparison, the equivalent differentiation regimen of C_2_C_12_ cells, as reported by Wang et al. (2019), is also provided. Peak differentiation was achieved after seven days, as evidenced by extensive structural rearrangement and the formation of polynucleated myotubes (Fig. 1B). RNA was isolated (n = 24) for 3’ end-focused RNA-seq library preparation (Quant-seq™) and single-end sequencing on the Illumina HiSeq 3000 platform. Raw sequencing data were processed using in-house bioinformatics pipelines for the statistical analysis of gene expression and alternative polyadenylation (Harrison et al. 2015).

**FIGURE 1.**
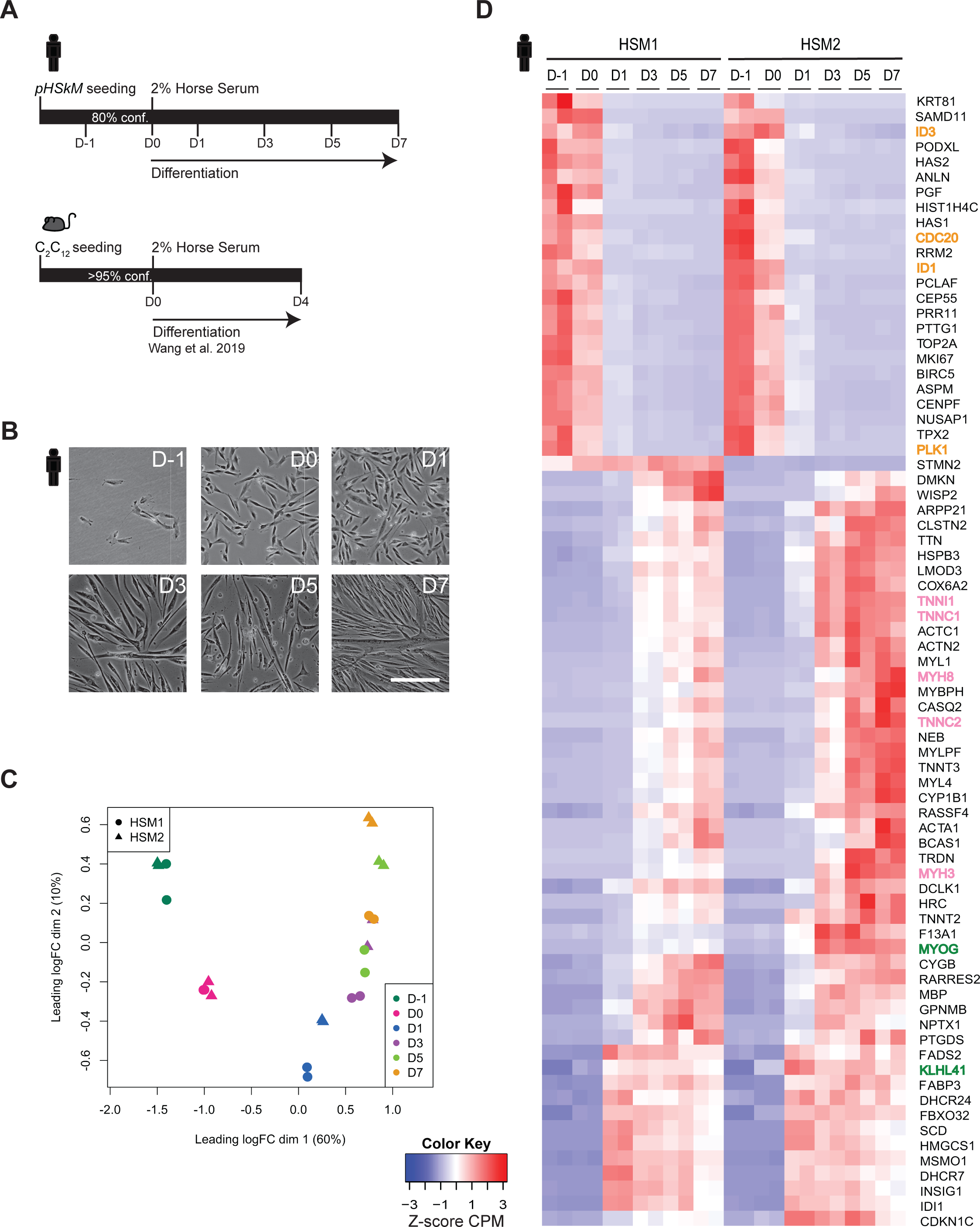
Reproducible differentiation of *pHSkM* cell lines. **(A)** Schematic representation of primary Human Skeletal Myoblast (*pHSkM*) differentiation (top) or C_2_C_12_ cells as reported by Wang et al. (2019) (bottom). Sample collection was at the time points indicated. **(B)** Grayscale brightfield microscopy images of *pHSkMs* at the indicated days of differentiation. The scale bar for all images is 100 μm. **(C)** MDS analysis of male-derived HSM1 (circle) and female-derived HSM2 (triangle) *pHSkMs*. The two replicates at each time point show discrete separation based on the differentiation of the myoblast lines (dim 1). For this analysis, the log2-normalised gene CPM were used for the top 2,500 genes. **(D)** Hierarchical clustering analysis of the top 75 significantly differentially expressed genes during *pHSkM* differentiation, illustrates overall reproducibility between the male-derived HSM1 and female-derived HSM2 cell lines (n = 2 replicates at each time point). Key gene categories, including downregulation of cell-cycle genes (orange), and upregulation of skeletal muscle contraction (pink) and muscle-specific differentiation markers (green) support efficacy of diff. The z-scores of gene CPM were clustered using Euclidean distance and single linkage method. Red is upregulated and blue is downregulated, as shown in the key.

Multidimensional scaling (MDS) demonstrated strong reproducibility between replicates and between donors (HSM1/2) along a shared trajectory of differentiation (Fig. 1C). As expected, *pHSkMs* and differentiated myotubes displayed distinct gene expression profiles, with the two *pHSkM* lines showing a high degree of replication despite intrinsic genetic and sex differences between the donors (Fig. 1D). The effectiveness of the differentiation protocol was supported by the downregulation of cell-cycle genes (e.g., *ID1*, *ID3*, *CDC20*, and *PLK1*, indicated in orange), the upregulation of key muscle-specific differentiation markers (e.g., *KLHL41* and *MYOG*, green), and genes associated with skeletal muscle contraction (e.g., *MYH3*, *MYH8*, *TNNC1*, *TNNC2*, and *TNNI1*, pink). Normalized data is available for interactive search and visualization here (https://degust.erc.monash.edu/degust/compare.html?code=fcdeaa05fc9e7e33af29b804fec86d37#) or as a list of all significantly regulated genes (Supplemental Table S1).

Of the 12,518 detected genes, only 295 genes (2.6%) were significantly differentially expressed (FDR < 0.05; Log2FC > 1) between the two donors on day 7 of differentiation (Supplemental Table S2). A search for enriched GO terms failed to identify any associated functional categories or obvious sex-linked differences.

Therefore, although further research into donor-specific differences could be of interest, the overall replication between donors here was used to justify their use as biological replicates in downstream analyses, for a total of four replicates at each time point.

### Primary Human skeletal muscle myoblasts undergo minor 3’ UTR lengthening

Previously, Wang et al. (2019) reported global lengthening of 3’ UTRs when comparing undifferentiated C_2_C_12_ myoblasts (D0) and after differentiation (D4) using the 3’ READS method (Wang et al. 2019). The C_2_C_12_ time points reflected 95% confluent myoblasts prior to induction of differentiation and differentiated myotubes, respectively (Fig. 1A). analogous comparison in *pHSkM* (D0 vs. D7). For a direct comparison and to minimize confounding effects from bioinformatic processing, the raw data from Wang et al. (2019) were reanalysed using ‘tail-tools’. Unlike what was observed in C_2_C_12_ cells, very little APA was seen when we compared our *pHSkM* data with previously published data. The only significantly shifted gene was *TPM2*, for which the apparent shift was due to elevated expression of an alternative terminal exon isoform (Fig. 2A). This represents an upregulation of an alternatively spliced isoform, rather than 3’ UTR switching *per se*, was confirmed by 3’ RACE (Fig. S1A).

**FIGURE 2.**
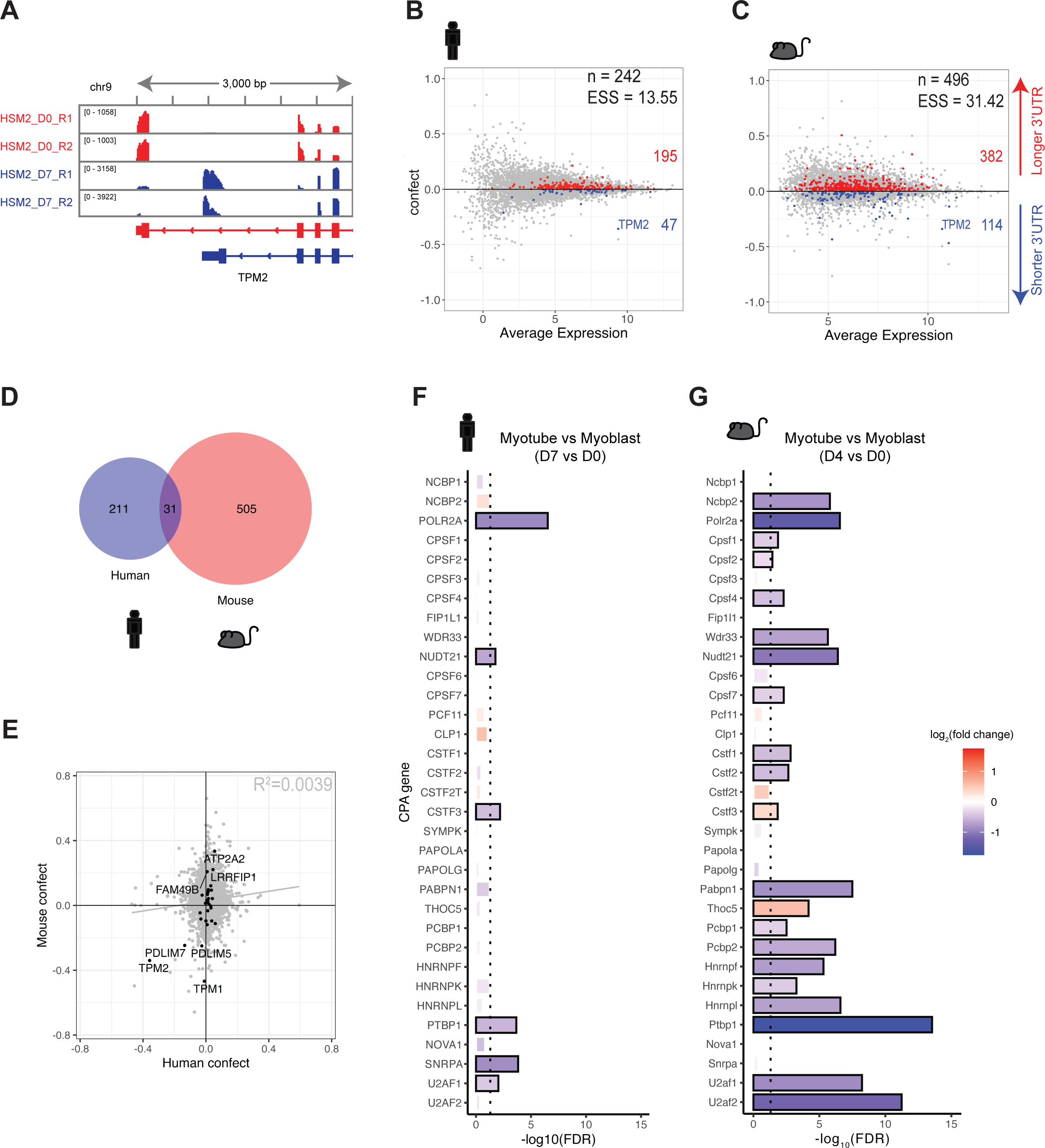
Differential CPA factor expression between models could explain APA differences. **(A)** IGV representation of reads mapping to *TPM2* 3’ UTR (Human version hg38). Proliferating *pHSkM* (HSM2 D0 R1 & R2) have reads (red) mapping to the distal poly(A) site of *TPM2*, while the primary myotubes (HSM2 D7 R1 & R2) have a greater number of reads (blue) mapping to the proximal poly(A) site. **(B-C)** Scatter plot of confect against average gene expression to visualise the shift in proportion of proximal versus distal poly(A) site usage. ESS (Estimated Sum of Squares) is an overall metric of the amount of APA, incorporating both the number of genes involved and the magnitude of shifting, intended to be broadly comparable across species and experiments (details in the Methods section). The degree of shift is quantitated by the estimated sum of squares (ESS) value. **(B)** *pHSkM* underwent less shift during differentiation (n = 242, ESS = 13.55) when compared to **(C)** C_2_C_12_ differentiation (n = 496, ESS = 31.42). Red represents 3’ UTR lengthening and blue represents 3’ UTR shortening. **(D)** Venn overlap of genes undergoing a shift in poly(A) site usage during *pHSkM* (D-1/D0 vs. D1/D3/D5/D7) and C_2_C_12_ (D0 vs. D4; n = 2 replicates at each time point) differentiation, with 31 genes in common. **(E)** Scatter plot of 31 overlapping shifting genes in *pHSkM* and C_2_C_12_ muscle differentiation. **(F)** Negative log10-normalised FDR of cleavage and polyadenylation (CPA) factors from (3). Myotubes vs. myoblasts, D0 vs. D7 in *pHSkM* and D0 vs. D4 in C_2_C_12_. FDR > 0.05 in dark grey. Nova1 expression was not detected in C_2_C_12_.

Because the RNA-seq methods used for both *pHSkM* and C_2_C_12_ data collection were 3’-focused, a side-by-side comparison was possible. Using raw data from Wang et al. (2019) reprocessed with tail tools (Harrison et al. 2015), a shift score for each gene in each sample was developed and implemented using the weighted matrix (weitrix) package (Harrison 2022). A rank-based score (between -1 and 1) was used to estimate the degree of APA between the groups of samples (myoblasts vs. myotubes). A “confect” score with a specified False Discovery Rate of 0.05 was used to rank genes by the magnitude of the APA shift (Harrison et al. 2019). To improve the chance of APA detection through increased statistical power, undifferentiated *pHSkMs* (D-1/D0) and differentiated myotubes (D1/D3/D5/D7) were pooled. Using this approach, 195 genes displayed 3’ UTR lengthening (positive shift) and 47 underwent 3’ UTR shortening upon *pHSkM* differentiation (Fig. 2B; Supplemental Table S3). In comparison, almost twice as many genes showed 3’ UTR lengthening (n = 382) or shortening (n = 114) in C_2_C_12_ cells (Fig. 2C; Supplemental Table S4).

Little correlation between gene expression change and APA was observed in either model, suggesting that changes in 3’ UTR usage occur independently of overall transcript levels (Supplemental Fig. S1B-C).

It is possible that subtle differences in methodology affect APA detection. Therefore, to quantify the global APA in an unbiased manner between unrelated experiments beyond the number of genes affected, a new method was developed. By the estimate of the sum of squares (ESS) approach, a numerical descriptor that considers the number of APA events, the degree of shift, and its variance can be calculated (see Methods). The ESS for APA in *pHSkM* was lower than that for C_2_C_12_ (ESS = 13.55 and 30.40, respectively), indicating that APA was significantly more pronounced in the murine model. By way of comparison, the knockdown of Pcf11, a core factor within the CPA with known impact on 3’ UTR choice (Sadowski et al. 2003; Ogorodnikov et al. 2018; Wang et al. 2019; Turner et al. 2020), displayed a high level of APA C_2_C_12_ cell line (ESS = 51.64) by re-analysis of (Wang et al. 2019) data (Supplemental Fig. S1D). Thus, using siPcf11 as a positive control, *pHSkM* and C_2_C_12_, underwent ∼26% and ∼59% APA, respectively.

To determine if the reduced APA in *pHSkM* was simply a less pronounced effect of the same genes, the human homologues of mouse APA genes were retrieved using Ensembl BioMart (38). Of the shared genes (n = 796) only 31 showed evidence of APA in both models (Fig. 2D). Of these, most shifts were in the same direction (Fig. 2E). However, the overall correlation was low (r^2^ = 0.0039), and clustering of the confect values close to the y-axis indicates that for most APA events, neither the genes nor their degree of shift was well conserved between *pHSkM* and C_2_C_12_ myotubes.

Previous studies have linked the differential expression of CPA factors to proliferative potential (Gruber et al. 2014). Highly proliferative cells, such as cancer cells, tend toward shorter 3’ UTRs (Sandberg et al. 2008) such that the 3’ UTR length is inversely proportional to the level of available CPA in most cell types where it has been measured (Sommerkamp et al. 2021; Turner et al. 2021). To determine whether the observed difference between APA in the two myotube models could be linked to the available CPA machinery, the expression of 33 previously identified CPA genes (Gruber et al. 2014) was analysed. In contrast to C_2_C_12_ cells, where most of the factors were differentially expressed, few of these showed significant differential expression in *pHSkM* (Fig. 2F-G). Collectively, these data support a model in which the paucity of APA associated with *pHSkM* differentiation compared to that observed in the immortalised C_2_C_12_ cells might be linked to the stability of CPA expression.

### Differentiating pHSkM and C_2_C_12_ myoblasts display major differences in gene expression

To understand if the immortalised murine C_2_C_12_ cell recapitulated the human differentiation processes more generally, we compared the gene expression profiles of the two models. The normalised gene expression data for C_2_C_12_ differentiation are available for interactive search here (https://degust.erc.monash.edu/degust/compare.html?code=c5d8df8af1b1dfe97477c254b30e876e#/) or as a list of all significantly regulated genes (Supplemental Table S5). A comparable *pHSkM* dataset included undifferentiated myoblasts (D0) and differentiated myotubes (D7). In total, 12,518 human genes were detected, 1,052 (8.5%) of which were significantly upregulated and 984 (7.8%) were significantly downregulated during differentiation (FDR < 0.05; Log2FC > 1) (Fig. 3A). In contrast, of 11,799 detected murine genes, 1,932 (16.4%) were significantly upregulated and 1,710 (14.5%) were significantly downregulated (FDR < 0.05; Log2FC > 1) (Fig. 3B). Interestingly, even with the statistical advantage of having twice the number of replicates (n = 4 replicates from 2 independent donors), *pHSkM* showed fewer differentially expressed genes and a lower magnitude of change than immortalised C_2_C_12_ cells (n = 2).

**FIGURE 3.**
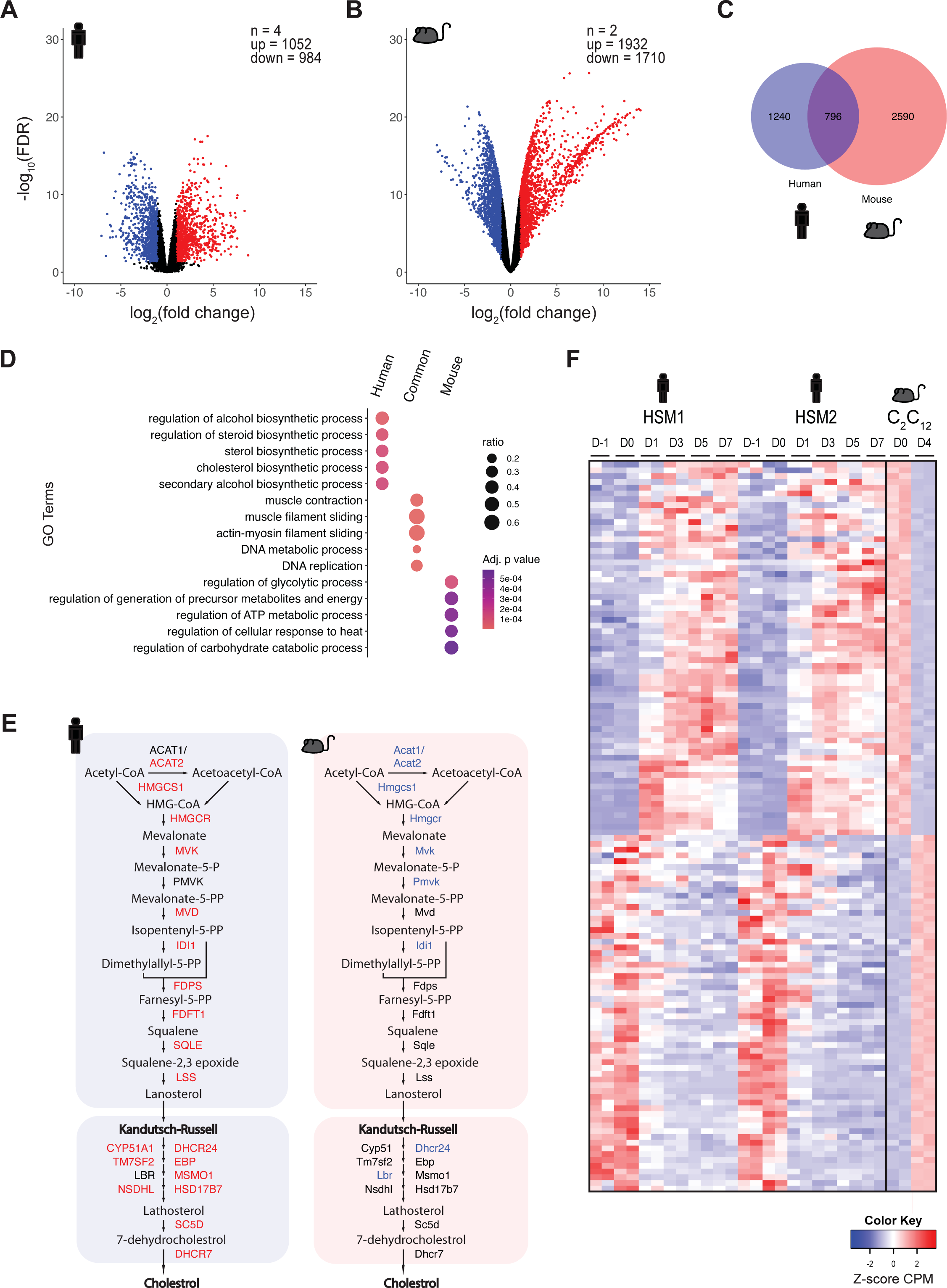
C_2_C_12_ and *pHSkM* differ on a molecular and metabolic level. **(A-B)** Volcano plot of statistical significance and fold change in proliferated versus differentiated *pHSkM* (D0 vs. D7; n = 4 replicates from 2 independent donors at each time point) **(A)** or C_2_C_12_ (D0 vs. D4; n = 2 replicates at each time point) **(B)** cells. Red indicates upregulated genes, and blue indicates downregulated genes (FDR < 0.05, Log2FC > 1). **(C)** Venn overlap of DEGs in *pHSkM* differentiation and human orthologs of DEGs in C_2_C_12_ differentiation, with 796 genes common. **(D)** Hierarchical clustering analysis of 796 overlapping genes during *pHSkM* and C_2_C_12_ differentiation. The z-scores of CPM were clustered using Euclidean distance and the single linkage method. Red signifies upregulated and blue downregulated, as shown in the key. **(E)** Gene Ontology (GO) analysis of DEGs specific to *pHSkM* (n = 1,240), common overlap (n = 796), and C_2_C_12_ (n = 2,590). The five most significant GO terms were then selected for each subset. Size and colour correspond to the gene ratio and adjusted p-value, respectively. With orange being the most significant. **(F)** Genes involved in cholesterol biosynthesis pathways (Mitsche et al. 2015; Meng et al. 2018). Red and blue indicate upregulated and downregulated genes, respectively. Black indicates no significant changes in gene expression during skeletal muscle differentiation.

To test for overlap between differentially expressed genes in the two models, human orthologs of the significant murine genes were retrieved using Ensembl BioMart (Smedley et al. 2009). Surprisingly, the overlap was modest. Of the 3,386 murine genes with shared human annotations, only 796 genes were differentially regulated in both datasets (Fig. 2C). To determine whether the lack of systematic replication between the two models had implications for biological function, GO associations were retrieved from the common and unique genes from the human or mouse lists using Enrichr (Chen et al. 2013; Kuleshov et al. 2016). The common subset was enriched, for processes specifically associated with muscle development, such as muscle contraction and actin-myosin sliding, as well as genes related to DNA processing, replication, and metabolic processes as expected, (Fig. 3D). Of the human-specific differentially expressed genes (n = 1,240), pathways related to alcohol, sterol, and cholesterol synthesis were enriched. A deeper analysis of the enzymes in the cholesterol biosynthetic pathway showed that most genes were upregulated in *pHSkM* (Fig. 3E). The complete gene set within the cholesterol biosynthesis GO term (GO:0006695) is shown (Supplemental Fig. S2A). This was not observed in mouse-specific differentially expressed genes (n = 2,590), where key genes in the cholesterol biosynthetic pathway were significantly downregulated. In contrast, differentiation in C_2_C_12_ cells was associated with the enrichment of genes involved in the GO term, regulation of glycolytic processes (GO:0006110) (Fig. 3D, Supplemental Fig. S2B).

Metabolic differences between myogenic models were recently highlighted in a meta-analysis comparing differences in microarray-based gene expression data for human, mouse, and rat myotubes, and corresponding tissue samples (Abdelmoez et al. 2020). Although not specifically noted in their analysis, the expression of *HMGCR*, the rate-limiting enzyme for cholesterol synthesis showed absolute expression difference between models when visualised using their convenient tool for interactive search https://nicopillon.com/tools/muscle-models-profiling/. Testing for statistical significance of normalised microarray-based expression from their raw data revealed that *HMGCR* and *DHCR24* were indeed expressed at a higher level in *pHSkM* than C_2_C_12_ myotubes, and more importantly, also in their respective primary tissue samples, indicating that the differences observed here were physiologically relevant (Supplemental Fig. S3; Supplemental Table S6).

In total, 130 orthologs were significantly regulated in the opposite direction between the models (Fig. 3F, Supplemental Table S7). Many such genes were supported by external data including, but not exclusive to processes such as cytokine signalling (*SOCS2*, *SOCS3* and *CCL2*), small molecule transport (*SLC43A3*), glycolysis (*GPD1*), lipogenesis (*PCSK9* and *FASN*) (Supplemental Fig. S3). Collectively, these results highlight that key metabolic pathways are differentially regulated between the *pHSkM* and C_2_C_12_ during differentiation.

## Discussion

An extensive literature links proliferation, exit from cell cycle, and differentiation to APA [reviewed by Manley (2017)]. Accordingly, differentiation in the immortalised C_2_C_12_ cell line, which is widely used to model muscle growth, function, and metabolism, has been linked to the widespread 3’ UTR lengthening (Li et al. 2015; Wang et al. 2019). Here we sought to study APA in Primary human skeletal myoblasts (*pHSkM*) differentiation, using established primary muscle cell lines derived from donor muscle biopsies (Zacharewicz et al. 2020). In this study, we show that unlike what has been observed in C_2_C_12_ myoblasts, differentiating *pHSkM* displayed only minor APA after differentiation. Moreover, transcriptomic analysis comparing *pHSkM* and C_2_C_12_ over a time course of differentiation revealed a substantial lack of conservation in transcriptional profiles as well.

For ease of reuse, transcriptomic changes in *pHSkM* (Fig. 1) and C_2_C_12_ differentiation (Fig. 3) can be accessed via an interactive data portal (https://degust.erc.monash.edu/degust/compare.html?code=fcdeaa05fc9e7e33af29b804fec86d37# and https://degust.erc.monash.edu/degust/compare.html?code=c5d8df8af1b1dfe97477c254b30e876e#/ respectively). The differences in enriched GO pathways between the two models were related to the major metabolic pathways (Fig. 3). In a meta-analysis of publicly available microarray data, Abdelmoez et al. (2020) reported that differentiated *pHSkM* and C_2_C_12_ myotubes employed different strategies for glucose metabolism (Abdelmoez et al. 2020). While both *in vitro* models favoured glycolysis, transcript levels suggest that C_2_C_12_ myotubes complete the oxidation of glucose through mitochondrial respiration, whereas *pHSkM* myotubes appear to favour a Warburg-like conversion of pyruvate to lactate. While both studies highlight metabolic differences, the Abdelmoez study and our results were derived from a fundamentally different experimental design. Abdelmoez et al. (2020), reported end-point differences in mRNA amount (normalised signal strength) in myotubes of different models as analysed by microarray (Abdelmoez et al. 2020). We report internal gene expression changes (myoblasts vs. myotubes) and subsequently compared the changes associated with differentiation between models.

Here, GO terms for carbohydrate metabolism were enriched in C_2_C_12_ compared to the *pHSkM* model, confirming that C_2_C_12_ more closely recapitulates the metabolic properties of fast muscle fibres. These glycolytic pathways were not enriched in differentiating *pHSkM*, suggesting they may display mixed metabolic properties. While C_2_C_12_ were originally established from mouse quadriceps muscle (Yaffe and Saxel 1977), whose white portion comprises 100% of type IIb muscle fibres (Campbell et al. 2001), *pHSkMs* were derived from skeletal muscle tissue samples biopsied from the *Vastus Lateralis* muscle, which includes a mix of oxidative and glycolytic fibres and up to 40% type I muscle fibres (Staron et al. 2000), with few sex-specific differences. Surprisingly, we observed a robust upregulation of alcohol, cholesterol, sterol, and steroid biosynthetic processes in *pHSkM* which were either unaffected or downregulated in C_2_C_12_ myotubes (Fig. 3E, Supplemental Fig. S3). Cholesterol is essential for myotube function, influencing electrical conduction, signalling, and membrane fluidity via raft-like structures in the T-tubule (Barrientos et al. 2017).

Cells are influenced by their genotype, including the complement of sex chromosomes present (Zore et al. 2018). Despite being derived from donors of each sex, the influence of sex chromosomes on gene expression in our *pHSkM* lines *in vitro* was limited. While muscle-derived cells retain some epigenomic memory in culture (Turner et al. 2020), the extent to which *pHSkM* lines maintain their identity, outside of the context of normal physiological control, needs further examination. The controlled environment of *pHSkM* cells in culture could reveal aspects of inter-individual and sex-specific muscle metabolism that are masked by the direct analysis of donor biopsies. However, such research would require a large number of donors to achieve statistical power. Collectively, our data highlight significant gene regulatory differences between the most common models used for *in vitro* study of muscle biology. Based on these differences we caution against their interchangeable use and suggest that, where it is feasible, the *pHSkM* model might more reliably model human muscle physiology.

The relatively minor APA associated with differentiation in *pHSkM versus* C_2_C_12_ cells was surprising. Multiple previous studies have connected myogenic differentiation to APA. Indeed, the expression of APA molecular drivers and the switch between proliferative and differentiated states has been well established. A simple explanation for the difference in APA is that unlike C_2_C_12_ cells, which display marked changes in CPA expression, *pHSkM* differentiation does not significantly alter their expression levels (Fig. 2F). However, APA and the expression of CPA factors have also been associated with proliferative potential (Gruber et al. 2014). Supporting a role for proliferative rate in contributing to differences in APA reported here, Abdelmoez et al. (2020), showed that C_2_C_12_ cells display as much as a 5-fold higher proliferative capacity than *pHSkMs* (Abdelmoez et al. 2020). However, in addition to legitimate differences in gene regulatory programs in the two models, we cannot rule out that contributions by experimental factors such as the number of proliferative cells remaining post-differentiation, and the transcriptomic contributions by any co-cultured non-myocyte cells such as fibroblasts in *pHSkM* are likely to impact our comparative analyses.

Finally, our data suggest that APA might be less marked in primary cell differentiation than has been reported by study of transformed cells. Recent work tracing APA in single-cell data (Agarwal et al. 2021; Wang et al. 2022; Zhou et al. 2022) brings a more fine-grained view of APA in native tissue than was previously possible from bulk RNA-seq analyses of tissues or cells in culture. For example, by extracting APA from single cells in cell-cycle resolved analyses (Wang et al. 2022), in a cell type-specific manner in the brain (Yang et al. 2021) and across murine during development, where a significant proportion (62%) of APA genes show longer 3’UTRs many (38%) do not (Agarwal et al. 2021). Furthermore, while some cell types, such as neuronal lineages show pronounced evidence of 3’UTR lengthening across developmental stages, others including myocytes do not (Agarwal et al. 2021). Thus, we believe that our findings fit into a broader narrative in which identification, and interpretation of the regulation and function APA will become more nuanced.

## Materials and Methods

### Primary human skeletal myoblast differentiation

The *pHSkMs* were obtained from two healthy human donors and established as previously reported (Zacharewicz et al. 2020) with ethical approval from the Deakin University Human Research Ethics Committee (2018-388). Isolated *pHSkMs* from the two donors, HSM1 (a 22.9 years old Caucasian male; 169 cm in height with a body mass of 63 kg) and HSM2 (a 26.5 years old Caucasian female; 176 cm in height with a body mass of 82.1 kg), were seeded in biological triplicates at 100,000 cells per well in 6-well plates with 2 ml of media. Cells were maintained and expanded in proliferation media, Ham’s F-10, with 20% Fetal Bovine Serum (FBS) and 2.5 ng/mL (0.01%) basic Fibroblast Growth Factor (bFGF). Once cells reached 80% confluence, the proliferation medium was replaced with high glucose Dulbecco’s Modified Eagle’s medium (DMEM) supplemented with 2% Horse Serum to initiate differentiation. All media were supplemented with 0.5% penicillin-streptomycin and 0.5% amphotericin B and refreshed every 48 h. The cells were maintained at 37°C and 5% CO_2_ in humid air. Brightfield microscope images were taken of the cells at each stage of differentiation at 10x magnification at a scale of 1.525 pixel/µm (TS100 DS-Fi2-L3, Nikon). Images were converted to grayscale without altering intensity or contrast.

### RNA extraction and sequencing

At each time point, two technical replicates were analysed per *pHSkM* line. For RNA sample collection, the medium was aspirated and 600 μl TRIzol™ was added directly to each 6-well plate well. After approximately 10 min of incubation at room temperature, a cell scraper was used to ensure that all cellular material was suspended. TRIzol™ extracts were stored at -80 °C until all samples were collected for parallel processing. To isolate RNA, samples were thawed on ice, and the RNA Directzol kit (Zymo) was used as per the manufacturer’s protocol.

Quant-seq, a 3’ end-focused RNA sequencing technique, was used to analyse global gene expression and alternative poly(A) site selection in *pHSkM*. The libraries were constructed using the QuantSeq 3’ mRNA-Seq Library Prep Kit FWD for Illumina (Lexogen). The libraries were prepared using 150 μg of input RNA according to the manufacturer’s instructions and sequenced at the Gandel Charitable Trust Sequencing Centre (Australia) using 150bp single-end reads on a single lane of the Illumina HiSeq 3000 instrument. The raw *pHSkM* data can be accessed from <u>GSE168897</u> (https://www.ncbi.nlm.nih.gov/geo/query/acc.cgi?acc=GSE168897). The 3’ end-focused 3’READS RNA sequencing data were retrieved from <u>GEO:GSE115232</u>(https://www.ncbi.nlm.nih.gov/geo/query/acc.cgi?acc=GSE115232) (17): C_2_C_12_ myoblast (GSM3190450, GSM3190451), C_2_C_12_ myotube (GSM3190452, GSM3190453), C_2_C_12_ siCtrl (GSM3171723, GSM3171724) and C_2_C_12_ siPcf11 (GSM3171727, GSM3171728).

### RNA-seq analysis

The sequencing data was aligned to the human genome (GRCh38, Ensembl release 93) or mouse genome (GRCm38, Ensembl release 93) using STAR aligner (Dobin et al. 2013), as implemented by the tail-tools pipeline (Harrison et al. 2015). To visualize the read coverage, we used the Broad Institute’s Integrative Genomics Viewer (IGV) (Robinson et al. 2011).

Following alignment, the differential gene expression analysis was conducted using Degust (Powell et al. 2019) using the Voom/Limma method (Law et al. 2014). To eliminate genes with low expression, we used a minimum gene read count cut-off of 50 in at least one replicate, along with a minimum gene CPM cut-off of two in at least three *pHSkM* or two C_2_C_12_ replicates. Genes exhibiting a fold change greater than two and a false discovery rate (FDR) < 0.05 were identified as statistically significant and considered to be differentially expressed. High-dimensionality reduction was achieved through Multidimensional Scaling (MDS) analysis using the ‘limma::plotMDS()‘ function (Ritchie et al. 2015) with the top 2,500 differentially expressed genes (DEGs).

In order to compare cross-species transcriptomic profiles, human orthologs of mouse genes were obtained from Ensembl Biomart (release 93) (Smedley et al. 2009). To identify genes with similar expression patterns, we executed hierarchical clustering on z-scores derived from gene CPM values. The clustering was conducted with the gplots package (version 3.1.0) (Warnes et al. 2020) using Euclidean distance measures and a single linkage approach. We used Enrichr for Gene Ontology analysis (Chen et al. 2013; Kuleshov et al. 2016) to identify enriched Biological Process terms associated with the gene sets.

To further reinforce the gene expression analysis, we examined the normalised gene counts from microarray data of human and mouse myotubes, along with corresponding adult skeletal muscle tissue. This data is accessible on GitHub at https://github.com/NicoPillon/Muscle_Models_Profiling/ (Abdelmoez et al. 2020). As the data was non-normally distributed for the genes considered in this study, we used the Kruskal-Wallis test with pairwise Dunn’s multiple comparisons to identify statistically significant differences in gene expression between species.

### Differential APA analysis using weitrix

Most APA detection methods do not control for variance well. Since samples with more reads provide better estimates of APA, these can be accounted for using observation-level weights similar to the limma-voom method (Law et al. 2014) for gene expression data. Here, a new method that improves on our previous APA detection (Turner et al. 2021) was implemented using the weitrix R package (Harrison 2022). For *pHSkM*, the expression of proliferating samples (D-1/D0, n = 8) and differentiated samples (D1/D3/D5/D7, n = 16) were compared. Similarly, for C_2_C_12_ the expression of proliferating samples (D0, n = 2) and differentiated samples (D4, n = 2) were compared. Meanwhile, for C_2_C_12_ siPcf11, the comparison focused on the Pcf11 knock-down (siPcf11, n = 2) and control (siCtrl, n = 2).

Specifically, we calculated summary scores of the APA state of each gene in each sample, compared to an average over all samples, with values between -1 (proximal) and 1 (distal). From these per-sample scores, we can estimate the change in APA state between the groups of samples. We also determine whether any such change is statistically significant and place an inner confidence bound on the change using the Topconfects method (Harrison et al. 2019). This method can be used for genes with two or more APA sites.

For each gene 𝑖, a set of 𝑛_𝑠𝑖𝑡𝑒 𝑖_ APA sites were identified. Let the read count in sample 𝑗 at site 𝑘 be 𝑐_𝑖,𝑗,𝑘_, the total count across samples be 𝑐_𝑖,⋅,𝑗_, across sites be 𝑐_𝑖,𝑗,⋅_ and across samples and sites be 𝑐_𝑖,⋅,⋅_. Assume that sites have been ordered from the furthest upstrand to the furthest downstrand. The rank-based score for all the reads at a site, 𝑠_𝑠𝑖𝑡𝑒 𝑖,𝑘_, was calculated using the proportion of reads upstrand of the site minus the proportion of reads downstrand of the site.

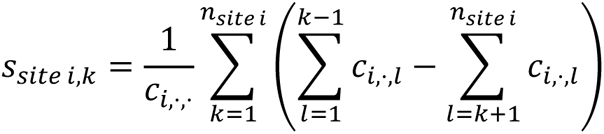

Then, for each gene 𝑖 and sample 𝑗, the score is calculated as the average of the site score of each read.

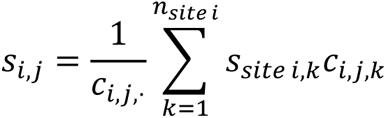

The variance of the average of 𝑐_𝑖,𝑗,⋅_ such reads will be 1/𝑐_𝑖,𝑗,⋅_ times the read-level variance. Different amounts of noise in particular observations are accounted for in linear models using observation-level weights that are inversely proportional to the variance. The appropriate weights are 𝑐_𝑖,𝑗,⋅_. If there were no reads for a particular sample in a particular gene, then the weight was zero, indicating missing data. Using these weights, an initial linear model was fitted to each gene.

If there is biological variation between individual samples, the variance will be somewhat higher than the suggested initial weights. If there are real differences between the groups of samples, the variance within each group will be somewhat lower. Therefore, we sought to refine the initial weights. We can also potentially use an estimate of the per-read variance for each gene under the assumption that the reads were observed at random.

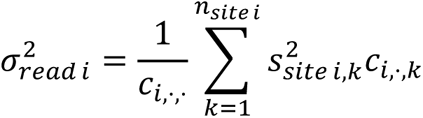

Calibrated weights were determined using residuals from the initial linear models fitted for each gene. A gamma GLM with a log link function of the squared residuals was fitted in terms of the log per-read variance 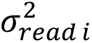 and a cubic spline curve of the log number of reads. This model was fitted to the entire dataset. The inverse of the predictions of this model provided a new set of weights. In other words, the new weights are calculated as a calibrated function of the number of reads and per-read variance.

Linear models were again fitted for each gene using the new weights. The final step of calibration is provided by Empirical Bayes squeezing of the residual variance for each gene, as in the limma. Shift effect sizes between groups are estimated as contrasts of linear model coefficients, as in the limma. The calibrated weights allowed accurate standard errors of these contrasts to be calculated.

### Quantifying APA with the Estimated Sum of Squares (ESS)

To quantify the extent of APA between independent samples beyond the raw number of genes, a new approach was required. A quantity that measures this is the sum of the squared APA shifts for all the genes. However, estimated shifts contain estimation errors that inflate the sum of squared shifts, and an adjustment for this is necessary.

For each gene, *i*, we regarded the estimated shift as a random variable 𝑆_𝑖_ composed of a true shift 𝑡_𝑖_ plus a random error 𝜀_𝑖_.

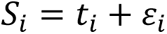

We seek an unbiased estimate of the sum of squares of the true shifts 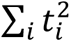. We assume 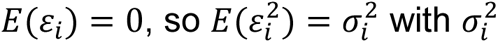 is the variance of the estimation error. The differential APA test provides an estimate of this variance as part of its output 𝑉_𝑖_, 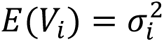.

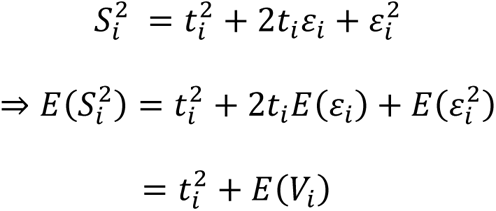

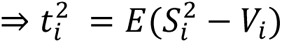

Thus, the square of the estimated shift minus the estimated variance is an unbiased estimate of 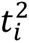, and summing these estimates over all genes provides an unbiased estimate of the sum of squares of the true shifts. Because this is adjusted for measurement accuracy, values are comparable between experiments performed with differing numbers of replicates, sequencing depth, and other sources of variation.

## 3’ RACE validation

The TVN-Poly(A) Test (TVN-PAT) was used to reverse transcribe polyadenylated RNA into cDNA for PCR-based 3’ RACE (Janicke et al. 2012). cDNA, prepared from 150 ng of total RNA input from each timepoint, was amplified with forward primers designed upstream of the proximal or distal poly(A) cleavage site: proximal forward primer (5’-GGAGCAGAAATTGCCAACAT-3’) and distal forward primer (5’-CGAGATTCACCAGACCTTGG-3’), along with a universal reverse primer (PAT reverse primer – 5’-GCGAGCTCCGCGGCCGCGTTTTTTTTTTTT-3’). For cDNA amplification, 5 μl of the cDNA, diluted at a ratio of 1:11, was PCR amplified using Amplitaq Gold 360 under the following conditions: 95°C for 10 min; 95°C for 20 s, 60°C for 20 s, 72°C for 30 s (27 cycles); and 72°C for 5 min. The amplified product was separated by TBE electrophoresis with 2% ultrapure agarose including SYBRsafe for imaging. Gels were imaged with Amersham^TM^ Imager 600. Linear uniform adjustments were made to image brightness and contrast.

## Data availability

The sequencing data produced in this study have been submitted to GEO under the accession number <u>GSE168897</u> (https://www.ncbi.nlm.nih.gov/geo/query/acc.cgi?acc=GSE168897).

## Supplemental Material

Supplemental Figures S1-S3

Supplemental Tables S1-S7

## Acknowledgements

We thank the members of the Beilharz and Lamon labs for their constructive feedback on the manuscript. The MHTP Gandel Medical Genomics Facility and Monash Bioinformatics Platform are acknowledged for providing technical and infrastructure support.

## Author contribution

AV performed bioinformatics and RNA analyses, PFH developed the weitrix and ESS bioinformatic packages for robust differential APA analysis, SEA isolated and differentiated donor *pHSkMs,* and AS performed RNA-seq and guided wet-lab RNA research. BD, SL, and THB conceived of and guided the study. AV, PFH, SL and THB wrote the manuscript. All authors provided feedback on the manuscript.

